# Anti-σ^28^ factor FlgM secretion regulates *Vibrio cholerae* adaptability in adult mice through quorum sensing and methionine metabolism

**DOI:** 10.64898/2026.01.02.697338

**Authors:** Guozhong Chen, Zixin Qin, Fenxia Fan, Mei Luo, Hongou Wang, Baoshuai Xue, Shucheng Li, Shiyong Chen, Xiaoman Yang, Xiang Mao, Liwen Yi, Chunrong Yi, Wei Li, Xiaoyun Liu, Biao Kan, Zhi Liu

**Affiliations:** Key Laboratory of Molecular Biophysics of the Ministry of Education/Department of Biotechnology, College of Life Science and Technology, Huazhong University of Science and Technology, Wuhan, Hubei 430074, China; State Key Laboratory of Microbial Technology, Shandong University, Qingdao, Shandong 266237, China; National Key Laboratory of Intelligent Tracking and Forecasting for Infectious Diseases, National Institute for Communicable Disease Control and Prevention, Chinese Center for Disease Control and Prevention, Beijing 102206, China; Department of Microbiology and Infectious Disease Center, NHC Key Laboratory of Medical Immunology, School of Basic Medical Sciences, Peking University Health Science Center, Beijing 100191, China; Key Laboratory of Zoonosis, Ministry of Education, College of Veterinary Medicine, Jilin University, Changchun, Jilin 130015, China; School of Marine Science and Engineering, Qingdao Agricultural University, Qingdao, Shandong 266109, China; Institute of Hygiene Toxicology, Wuhan Centre for Disease Prevention and Control, Wuhan, Hubei 430024, China; Department of Infectious Diseases, Peking University Third Hospital, Beijing 100191, China

**Keywords:** *Vibrio cholerae*, flagellum, FlgM, quorum sensing, methionine metabolism

## Abstract

**Introduction:** Cholera, caused by *Vibrio cholerae*, is a severe diarrheal disease threatening global health. The flagellum of *V. cholerae* functions not only for motility but also as an environmental sensor regulating virulence. The trade-off mechanism between motility loss and enhanced host adaptation remains unclear.

**Objective:** This study aimed to elucidate how flagellar mutations, which lead to a loss of motility, affect host adaptability in *V. cholerae* and to uncover the underlying molecular basis.

**Methods:** We analyzed 3,135 cholera-related samples to identify mutation hotspots in flagellar genes. A relevant flagellar mutant library was constructed and assessed for host adaptability. Multiple approaches, including molecular, genetic, transcriptomic, and proteomic analyses, were applied to dissect the FlgM-mediated pathway, with key findings revalidated in an adult mouse model using a specifically constructed 6N-labeled mutant pool.

**Results:** Big data analysis reveals those flagellar mutations in *V. cholerae* cluster in key structural and regulatory genes, including *flhB*, *fliA*, *fliF*, *fliD*, and *fliM*. Flagellar mutations led to a scenario where the secretion level of the regulator FlgM was negatively correlated with host adaptability. Intracellular FlgM inactivates either the σ^28^ factor FliA directly or acts through the VarS/VarA-CsrA/BCD system, ultimately leading to the derepression of the quorum-sensing master regulator named *hapR*. HapR can directly bind to the promoters of genes involving in methionine transportation to regulate adaptability of *V. cholerae*. The 6N-labeled mutants pool experiment reconfirmed that motility loss promotes host adaptability via the FlgM-FliA-HapR-methionine axis.

**Conclusion:** Our findings demonstrate that secretion levels of the flagellar regulator FlgM drive an adaptive shift in *V. cholerae* from motility to host adaptability, mediated through quorum sensing and methionine metabolic reprogramming. This reveals a novel mechanism underlying bacterial evolution and pathogenicity.

## Introduction

Cholera, caused by *Vibrio cholerae*, is characterized by severe watery diarrhea that can be lethal. Infection occurs through the consumption of contaminated food or water. Since 1817, there have been seven global pandemics that resulted in massive human deaths, substantial economic losses and considerable healthcare burdens [1]. The WHO reports a large number of cholera cases or outbreaks, from 44 countries in 2022 to 45 countries in 2023, which poses a serious challenge of cholera prevention and control [2].

The life cycle of *V. cholerae* is closely linked to its pathogenic mechanism. Initially, the bacteria enter the host via contaminated water, survive gastric acid, and colonize the small intestine. Subsequently, their flagella facilitate penetration of the mucus layer, allowing access to the intestinal epithelial surface. As infection progresses, *V. cholerae* forms biofilms and employs quorum-sensing regulation. Upon reaching high cell density, the bacteria detach from the epithelial surface, enter the intestinal lumen, and are ultimately excreted in host feces. They then return to aquatic environments, thereby completing the transmission cycle [3, 4].

*V. cholerae* is a Gram-negative bacterium with a single polar flagellum, by which it swims rapidly and this motility contributes to quick transmission among different habitats [5,6]. As the propeller of *V. cholerae* movement, the flagellum is a sophisticated device composed of many different structural and functional components. Its biosynthesis involves expression of proteins involved in a series of fine and accurate regulations, acting on transcriptional, translational, and post-translational levels [7]. Transcriptional regulation of *V. cholerae* genes has been well-studied, and is built upon a four-tiered transcriptional hierarchy [3]. The flagellum of *V. cholerae* is also involved in many other biological activities, such as promoting adherence [8], influencing the infectivity [9], triggering c-di-GMP production [10], and sensing mucus layer mechanical attraction [4]. Interestingly, a high number of single nucleotide polymorphisms (SNPs) were detected in the Haiti and Nepal isolates of the 7^th^ cholera pandemic, with 7 out of 54 high quality SNPs belonging to flagellum-associated genes [11]. Similarly, a large number of mutants with impaired motility have been isolated from the laboratory [12–17], the natural environment [18–20], and clinical samples [21]. However, the relationship between motility loss and host adaptability in *V. cholerae* remains unclear.

Here, we analyzed 3,135 cholera-related samples and identified a recurrent pattern of flagellar mutations in *V. cholerae*, with notable clustering in genes including *flhB*, *fliA*, *fliF*, *fliD*, and *fliM*. Screening a flagellar mutant library revealed an inverse correlation between bacterial colonization capacity and FlgM secretion. Mechanistically, elevated intracellular FlgM inhibits the σ^28^ factor FliA, which derepresses transcription of the quorum-sensing master regulator *hapR*. Subsequent upregulation of methionine metabolism and transport by HapR enhances *V. cholerae* colonization in vivo. Using a uniquely constructed 6N-labeled mutant pool in an adult mouse model, we further confirmed that loss of motility promotes host adaptability through the FlgM-FliA-HapR-methionine regulatory axis. Together, our data will shed light on trade-off mechanisms between motility loss and host adaptability.

## Materials and methods

### In silico analysis flagellar gene mutations in V. cholerae

A total of 3,135 samples in Table S1 were retrieved from the NCBI SRA database by searching for “cholera” or “Vibrio cholerae” in combination with metagenomics-related keywords. These valid sequencing runs were selected for subsequent analysis of flagellar gene mutations. For each dataset, raw reads were quality-filtered using fastp v0.23.2 [22] and assembled with metaWRAP v1.3.2 [23]. Resulting contigs were binned and taxonomically classified using GTDB-Tk v2.4.0 [24]. Bins identified as *V. cholerae* were retained for protein prediction with Prodigal v1.2.6 [25]. A custom reference library of 51 flagellar proteins in Table S2 was constructed based on the *V. cholerae* N16961 genome. Flagellar mutations across 1258 assembled *V. cholerae* bins were then identified in Table S3 and quantified using DIAMOND BLASTP v0.9.30-3 [26]. As controls, the *hapR* gene (which carries a frameshift mutation in N16961) served as a positive control, while the housekeeping genes *asd* (VC2107), *mdh* (VC0432), and *dnaE* (VC2245) were used as negative controls, respectively [27,28]. The codes for 6N-Label relative abundance analysis, bins acquisition and species annotation from cohort-based studies, and analyzing high-frequency mutations in the flagellar genes of *V. cholerae* of this study are openly available in Zenodo at https://doi.org/10.5281/zenodo.14920964, reference number 14920964.

### Bacterial strains, plasmids, and growth conditions

All *V. cholerae* mutants were derived from strain E1 Tor C6706 and cultured in Luria-Bertani (LB) medium (Qingdao Hope Bio-technology, China), M9 minimal medium containing 2 mM MgSO_4_, 0.1 mM CaCl_2_ and 0.2% casein acid hydrolysate (Sangon Biotech (Shanghai) Co., Ltd., China), or defined medium containing appropriate antibiotics (streptomycin 100 μg mL^-1^, kanamycin 50 μg mL^-1^, ampicillin 100 μg mL^-1^, chloramphenicol 25 μg mL^-1^ for *E. coli* and 1.0 μg mL^-1^ for *V. cholerae*) or 1mM isopropyl β-D-1-thiogalactopyranoside (IPTG) at 37 °C unless otherwise noted. Antibiotic and IPTG powders were purchased from Sangon Biotech (Shanghai) Co., Ltd.. This defined medium was prepared with the omission of Met and supplementation with 0.5% glucose (Sangon Biotech (Shanghai) Co., Ltd., China) [29]. In-frame deletion mutants were constructed by cloning the regions flanking a target gene into suicide vector pWM91 that contains a *sacB* counter-selectable marker [30]. Complementation of *flgM* was achieved by inserting P*_lac_*-*flgM* into chromosomal *lacZ* using the suicide plasmid pJL1 [31]. Overexpression of *flgM* was obtained by cloning *flgM-his_6_* downstream of the *tac* promoter of pMal-c2x. Plasmid pBAD24 containing a pBAD-inducible promoter was used to overexpress *flgM-his_6_*, *flhB*, *flaA-his_6_*, *fliA-flag*, and *hapR-flag*. The fragment of pBAD-*fliA-flag* was cloned into pYM24 derived from pACYC117 [32], achieving coexistence with the plasmids pMal-*flgM-his_6_* and pBBR-*hapR-luxCDABE*. All constructed mutants and plasmids were confirmed by sequencing. All bacterial strains, primers and plasmids used in this study are listed in Table S4&5.

### Motility assay

*V. cholerae* strains were incubated overnight at 37 °C for 12 h. Single colonies were inoculated with sterile toothpicks into LB plates containing 0.3% select agar (Invitrogen, USA), and these soft agar plates were incubated at 37 °C for 8 h. Photographs were taken and analyzed for colony width as a measure of motility using a gel imaging instrument (GenoSens 1860, China).

### H_2_O_2_ killing experiments in vitro

Overnight cultures of flagellar mutants were 1:1,000 sub-cultured into M9 minimal medium containing 2 mM MgSO_4_, 0.1 mM CaCl_2_ and 0.2% casein acid hydrolysate and grown at 37°C, 200 rpm until mid-log phase. H_2_O_2_ (Sinopharm Chemical Reagent Co., Ltd., China) was directly added into the cultures at appropriate concentrations, followed by stationary incubation at 37°C for 30 min. Serial dilutions of the samples were plated on LB agar for enumeration of colony-forming units (CFU) and the survival rate in presence of H_2_O_2_ was calculated by comparing CFUs of H_2_O_2_-challenged and unchallenged samples [32].

### Biofilm formation

Colonies grown on LB agar plates were resuspended in LB broth and grown to an optical density at 600 nm (OD_600_) of ∼0.6. A 1:100 dilution was inoculated into 10 × 75 mm borosilicate glass tubes containing 1 mL M9 minimal medium with 2 mM MgSO_4_, 0.1 mM CaCl_2_ and 0.2% casein acid hydrolysate. Biofilms were allowed to form by static incubation at 37 °C for 96 h. The biofilms were stained with crystal violet (Solarbio Life Sciences Co., Ltd., China) and resolved with DMSO (Sinopharm Chemical Reagent Co., Ltd., China), quantifying with a microplate reader (Spark, Tecan, Switzerland) at 570 nm.

### In vivo colonization assays

The streptomycin-treated adult mouse model was used to assess the competition ability of mutants or complement strains during co-colonization with C6706 WT. ROS resistance in the adult mice assay was performed as previously described [33]. For this, five-week-old CD-1 mice (Jiangsu GemPharmatech Co., Ltd., China) were provided with drinking water with or without 10 g L^-1^ of the antioxidant N-acetylcysteine (MERCK, Germany) for one week. Afterwards, 5 g L^-1^ streptomycin and 0.05 g L^-1^ aspartame (Macklin Inc., China) were added to the drinking water. One day after streptomycin treatment, the mice were administrated with 100 μL of 10% (w/v) NaHCO_3_ (Sangon Biotech (Shanghai) Co., Ltd., China) followed by 100 μL of a 1:1 mixture of WT and mutant (approximately 10^9^ CFU for each strain per mouse). Fecal pellets were collected from each mouse at 1, 3, and 5 days after gavage, homogenized, serially diluted in LB and plated on LB agar containing streptomycin and 5-bromo-4-chloro-3-indolyl-β-D-galactopyranoside (X-Gal) (Sangon Biotech (Shanghai) Co., Ltd., China) to quantify the bacterial loads. The competition index CI was calculated as the ratio of mutant to WT colonies normalized with the input ratio. Five-day-old CD-1 mice were transferred to an incubator and left to adapt for 2 h at 30 °C before inoculation. The mice were then administrated with a 50 μL of a 1:1 mixture of WT and mutant *V. cholerae* (approximately 10^6^ CFU for each strain per mouse) and returned to the 30 °C incubator. The animals were sacrificed 18 h after gavage. Samples of the small intestine were removed and homogenized in 1.5 ml of LB, serially diluted, and plated on LB agar containing streptomycin and X-Gal for quantification of bacterial load. The CI was calculated as the ratio of mutant to WT colonies normalized with the input ratio.

### Growth curve determination and luminescence detection

Overnight LB cultures of targeted strains (with or without plasmid) were inoculated in fresh medium at a ratio of 1:100 and grown with shaking to mid-log phase at 37 °C. The bacteria were harvested by centrifugation, washed three times and resuspended to an OD_600_ of 1.0 with fresh pre-warmed medium (LB or M9), and then diluted 1:1,000 in 96-well microplates that were incubated at 37 °C under shaking. The growth kinetics were determined by measuring OD_600_ or OD_655_ using a microplate reader (Spark, Tecan, Switzerland) combined (where applicable) with luminescence detection. If necessary, 0.1 mM isopropyl-β-D-thiogalactoside (IPTG) or 0.2% arabinose was added. Readouts were performed every 30 min for 20 h. The luminescence assay was standardized at OD_600_. All experiments were performed in triplicate.

### Transmission electron microscopy (TEM) imaging

Single colonies of target strains were inoculated into 50 μL of saline and gently mixed with a clipped, sterile pipette tip. Five microliters of sample were added to a copper supporting mesh. Excess liquid was removed after standing in a ventilated and clean environment for 5 min. The mesh was stained with freshly prepared 1% phosphotungstic acid (Sangon Biotech (Shanghai) Co., Ltd., China) for 2 min. Excessive dye was removed and the mesh was dried for approximately 1 h prior to transmission electron microscopy (HT7700, HITACHI, Japan).

### Secretion of FlgM

To investigate the secretion of FlgM, culture supernatant of the *V. cholerae* mutants containing pBAD24-*flgM-his_6_* was collected and the protein was detected according to a previous method with slight modification [4]. All mutant sediment samples were standardized with CFU (10^9^). Proteins in the 400mL supernatant were concentrated with 20% trichloroacetic acid (TCA) (Sinopharm Chemical Reagent Co., Ltd., China). These, together with the proteins in the precipitate and supernatant were separated by SDS-PAGE, transferred to a polyvinylidene fluoride (PVDF) membrane (Millipore, USA). The proteins on the Western blot were detected with primary 6× His-tagged rabbit antibodies (Sangon Biotech (Shanghai) Co., Ltd., China) at a dilution of 1:5,000, and as secondary antibody horseradish peroxidase [HRP]-conjugated goat anti-rabbit immunoglobulin G (Sangon Biotech (Shanghai) Co., Ltd., China) was used at a dilution of 1:10,000. Immunoblotting was performed using the Pierce ECL Western Blotting Substrate Kit (Thermo Scientific, USA).

### Proteomic analysis

Overnight LB cultures were inoculated 1:100 in fresh LB medium and incubated until OD_600_ reached 0.9 (approximately 10^9^ CFU per sample). Then the culture was centrifuged. The pellets were for bacterial proteome samples. For the analysis of the bacterial secretome, the 400 mL of culture supernatant was filtered through a 0.22 μm microporous membrane to remove intact bacteria. The proteins in the filtrate were precipitated overnight at 4 °C with 20% TCA containing 0.5 mM sodium deoxycholate. After centrifugation at 10,000 g for 1 h at 4 °C, the protein pellets were washed three times with cold acetone for LC-MS/MS experiments, that were carried out as previously described [32]. The secretome experiments were performed in duplicates, and bacterial proteome samples were analyzed in triplicates. Proteins with Fold changes >1.5 and *P*< 0.05 were considered as significant differences.

### Gene expression analysis via RT-qPCR

Overnight LB cultures were inoculated 1:100 in fresh LB medium containing appropriate antibiotics or, if needed, 0.2% arabinose, and incubated at 37 °C. RNA for real-time PCR (RT-PCR) was extracted using an RNA extraction kit (Vazyme Biotech Co., Ltd. China). The RT-PCR reaction was performed using SYBR green fluorescent dye (Yeasen Biotechnology (Shanghai) Co., Ltd., China) and CFX Connect Real-time Detection System (Bio-Rad, USA). The genes *recA* or *secA* or 16S rRNA were used for normalization. Relative expression levels were calculated using the 2^-ΔΔCT^ method. All experiments were performed in triplicate. For reverse transcription, approximately 400 ng of RNA was extracted using the HiScript II 1st Strand cDNA Synthesis Kit (Yeasen Biotechnology (Shanghai) Co., Ltd., China). The RT-qPCR primers used for detection of mRNA of specific genes are listed in Table S5. All primers were tested for PCR specificity and efficiency.

### Protein purification

The proteins HapR-His_6_ were expressed in *E. coli* BL21 (DE3) and purified using the pET system and BeyoGold™ His-tag Purification Resin (Beyotime, China).

#### EMSA

The purified recombinant proteins HapR containing 6×His-tags were used for EMSA. The promoters of their target genes were amplified by PCR and purified. These amplicons (50 ng) were incubated with 6× His-tagged protein in 20 μL containing 4μL of 5× binding buffer (50 mM Tris-HCl, 0.25 M KCl, 2.5 mM DTT, 5 mM MgCl_2_, 0.25 mg mL^-1^ BSA, 2.5 mM EDTA, 1% glycerol, pH 8.3) (Sangon Biotech (Shanghai) Co., Ltd., China) at 25 °C for 30 min. DNA-protein complexes were analyzed by electrophoresis on a 5% native polyacrylamide gel at 4 °C and 80 V cm^-1^ for 1.5 h. After staining the gel in ethidium bromide for 10 min, a gel imaging system (GenoSens 1860, China) was used to produce images. The EMSA primers used in this study are listed in Table S5.

### Synthesis of fluorescent monomers

FITC-D-Glucose was synthesized as previously described [34]. Briefly, D-glucosamine hydrochloride (77.6 mg, 0.36 mmol, 1.2 equiv) and Et3N (91.1 mg, 0.9 mmol, 3.0 equiv) were mixed in DMF (10 mL) (Sinopharm Chemical Reagent Co., Ltd., China). After 1 h of stirring, fluorescein (116.8 mg, 0.3 mmol, 1.0 equiv) was added. The reaction mixture was stirred for 16 h at RT in the dark, after which the solid product was obtained by filtering and concentrated *in vacuo*.

### Bacterial metabolic activity assay in vitro

Bacterial cultures in LB were harvested at OD_600_ ∼0.4 by centrifugation and the cell pellet was washed 3 times with Met-free MM medium. The cells were resuspended and incubated at 37□ for 30 minutes. A solution of 10 mL Met-free MM medium and 1 µL of FITC-D-Glu was added and 500 µL of this was added to transwell wells and 200 µL of this was added to the transwell chambers. To each well, 100 µL of bacterial suspension was added and the plates were incubated for 30 min at 37 °C in the dark. The supernatant was washed off with PBS buffer and then assayed by Flow Cytometry (Agilent Technologies Inc., USA).

### Bacterial metabolic activity assay in vivo

Bacterial metabolic activity was assayed in vivo as previously described with following modifications [35]. Briefly, the CD-1 mice were administered with WT and mutants for competition colonization experiments as described above. At 48 h post-inoculation, the mice were administered with 100 μL FITC-D-Glu (0.1 mM) in PBS (Sangon Biotech (Shanghai) Co., Ltd., China). Two hours after the gavage, the animals were sacrificed and the small intestine was dissected and rinsed with 2 mL PBS, cut longitudinally and the intestinal content was scraped off using a cell scraper. The intestinal content was suspended into a 1.5 mL PBS buffer with 2 steel beads and shaken to release bacteria. Centrifugation at 3000 rpm for 1 min removed large tissue fragments and impurities, and the supernatant was filtered through 40 μm cell strainers to remove smaller cell debris. The filtrate was centrifuged (8,000 rpm, 5 min), the bacterial pellets were washed twice with 1.5 mL PBS and resuspended in PBS for flow cytometry analysis. In the FITC channel, cells with strong signals of approximately 10% were selected for sorting, with 10^5^ events obtained for each sample, and the collected samples were diluted and plated on LB agar containing streptomycin and X-Gal for quantification of the bacterial index.

### Detection of amino acid profiles in ΔflhB and ΔflaA by HPLC and MS

Overnight LB cultures of Δ*flaA* and Δ*flhB* were sub-cultured with an inoculum of 1:100 in MM medium and shaken at 37 °C until OD_600_=0.5. The bacteria were collected by centrifugation (>200mg) and samples of 60 mg were suspended in 50 μL water, vortexed for 60 s, and mixed with 400 μL methanol-acetonitrile solution (1:1, v/v) and 50 μL of a mixture of 16 isotopic internal standards (50 μM). The mixture was vortexed for 60 s and twice subjected to low-temperature ultrasound for 30 min. Proteins were precipitated at -20 °C for 1 h and after centrifugation at 14,000 g and 4 °C for 20 min and the supernatant was collected, freeze-dried, and stored at -80 °C prior to high-performance liquid chromatography (HPLC). HPLC was performed using an Agilent 1290 Infinity LC ultra-high-performance liquid chromatography system (USA). The mobile phase consisted of 25 mM ammonium formate + 0.08% formic acid (solution A) and 0.1% formic acid acetonitrile (solution B). The samples were loaded into an automatic sampler at 4 °C, and the column temperature was set at 40 °C. The flow rate was 250 μL min^-1^, and the injection volume was 1 μL. The gradient elution was as follows: 0-12 min, linear change from 90% B to 70% B; 12-18 min, linear change from 70% B to 50% B; 18-25 min, linear change from 50% B to 40% B; 30-30.1 min, linear change from 40% B to 90% B; 30.1-37 min, maintained at 90% B. At regular intervals between experimental samples a quality control (QC) sample was inserted to evaluate the stability and reproducibility of the system.

Mass spectrometry (MS) analysis was performed using a 5500 QTRAP mass spectrometer (SCIEX) in positive ion mode. The ESI source conditions for the 5500 QTRAP were as follows: source temperature 500 °C, ion Source Gas 1 (Gas 1): 40, Ion Source Gas 2 (Gas 2): 40, Curtain gas (CUR): 30, IonSapary Voltage Floating (ISVF) 5,500 V. The analytes were detected using the MRM mode.

### In vivo colonization assay of 6N-labeled mutants

Briefly, using C6706 (*vc1807*:: CmR) as a template, amplify the CmR fragment and insert a 6N site as a marker in the middle of one end primer (6N-R) [36], then clone it into the pWM91 vector. Sequence tags of 6N were inserted into the *lacZ* position of the 26 mutants and the WT strain with the 6N-labeled plasmids pWM91-CmR-6N by conjugation experiments. After labeling the bacterial strains, confirmed by sequencing, they were sub-cultured and adjusted to an OD_600_ of 1.0. After harvesting by centrifugation, the bacterial pellets were stored at -20°C. Using the methionine-supplemented and water-restricted mouse model, the mice were gavaged with 100 μL of the mixed bacterial samples per mouse. On day 5 after gavage, mouse feces were collected and total DNA was extracted, along with the input samples. Amplicons produced by PCR with primers targeting both ends of *LacZ* were sequenced. The sequenced raw data was analyzed by Vsearch for merging paired-end reads and Python to calculate the proportion of reads from the 6N-labeled mutants among the total reads.

## Statistical analysis

Results are presented as mean ± standard deviation. When two groups were compared, *t*-tests were performed. One-way analysis of variance (ANOVA) was used for two or more treatment groups. Statistical analysis was performed using the software GraphPad Prism v.9 for Windows. Differences with a p value less than 0.05 were considered to be statistically significant.

## Results

### High-frequency mutations in the flagellar genes of V. cholerae

The life cycle of *V. cholerae* primarily alternates between aquatic environments and the human host [3]. To investigate its adaptation across these niches, we categorized cholera-related metagenomic sequencing samples from the SRA database in Table S1 into five groups based on their origin: human, clinical, aquatic (including water and sewage), fish, and amoeba-derived. These samples were collected from 19 countries (Fig. 1A). A total of 1,258 metagenome-assembled genomes (MAGs) of *V. cholerae* in Table S3 were reconstructed from these samples. Their flagellar protein mutations were then systematically analyzed using the 51 flagellar proteins of *V. cholerae* N16961 in Table S2 as reference amino acid sequences. For quality control, the mutation-prone *hapR* gene, which carries a frameshift mutation in the N16961 genome, served as a positive control, while the conserved housekeeping genes *asd* (VC2107), *mdh* (VC0432), and *dnaE* (VC2245) were used as negative controls, as established in prior studies [27,28].

**Fig. 1.**
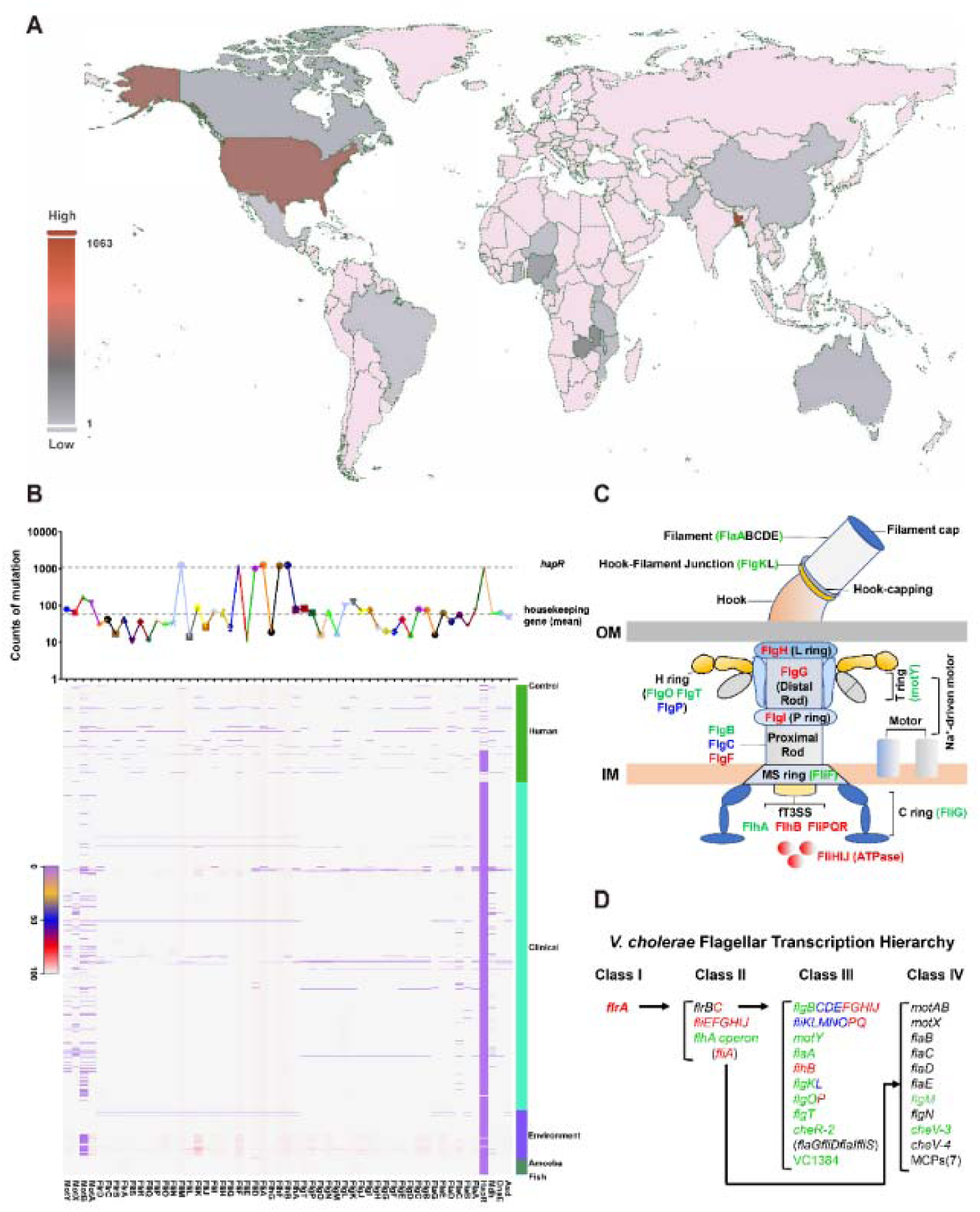
High-frequency mutations in the flagellar genes of *V. cholerae* from the life cycle. (**A**), World distribution map of the collected samples, with areas in pink indicating no sample collection. (**B**) Spectrum and count statistics of flagellar gene mutations. The *hapR* and housekeeping genes (*asd*, *dnaE*, *mdh*) were used as controls. (**C**) Schematic diagram of the *V. cholerae* flagellum assembly. (**D**) *V. cholerae* flagellar transcription hierarchy.

Our mutation frequency analysis revealed that across all sampled habitats, flagellar gene mutations in *V. cholerae* consistently clustered in key genes, including *flhB*, *fliA*, *fliF*, *fliD*, and *fliM* (Fig. 1B). Notably, mutations were also detected in the primary flagellar regulator gene *flrA*, a finding consistent with previous reports [16,37]. Functionally, these mutated genes are involved in critical flagellar structures and regulatory hierarchies, including Class I and IV transcriptional regulation, the type III secretion system (T3SS) apparatus, the filament-capping complex, and the C-ring switch structure (Figure 1C and D). This indicates that these loci have key evolutionary and pathological significance in regulating the host adaptability balance.

### Phenotypic screening of the flagellar mutant library

To elucidate the environmental adaptability conferred by the flagellar gene mutations identified through the above big data-driven analysis, a systematic mutant library targeting 26 flagellar-related regulons of *V. cholerae* listed in Table S4 was constructed using the primes in Table S5. Subsequently, this library was subjected to comprehensive phenotypic screening, assessing motility, biofilm formation, growth rates in both rich and minimal media, colonization efficiency in infant and adult mice, as well as resistance to HCOC. The HCOC assay is particularly relevant as it reflects bacterial survival under host-derived reactive oxygen species (ROS) stress. Of the investigated mutants, 22 mutants had completely lost their motility. Deletion of chemotaxis-related genes *cheV-3*, *flgO* and *flgP* resulted in significantly impaired motility (Fig. 2A, Fig. S1), which is consistent with previous reports [6, 38]. The colonization ability in infant mice was seriously damaged (Fig. S2) and biofilm formation of the 26 mutants were mostly enhanced, which were consistent with earlier findings (Fig. S3) [5,6,10]. Surprisingly, we observed completely opposite results in the adult mice, as some of the mutants were able to outcompete the WT strain (Fig. 2B), although their growth rates were consistently lower than that of WT in minimum or rich medium (Fig. S4 and 5), and others were attenuated. In total, five mutants (Δ*flgBCDE*, Δ*fliK*, Δ*fliKLMNOPQRflhB*, Δ*fliE,* and Δ*flgP*) produced no significance difference in CI (blue in Fig. 1D), nine mutants (Δ*flrA*, Δ*rpoN*, Δ*flrC*, Δ*fliEFG*, Δ*fliHIJ,* Δ*fliA*, Δ*flgFGHIJ,* Δ*fliPQR*, and Δ*flhB*) displayed enhanced CI (red in Fig. 2B), and twelve mutants (Δ*flhA*, Δ*flhA* operon, Δ*flgB*, Δ*motY*, Δ*flaA*, Δ*flgK*, Δ*flgO*, Δ*flgT*, Δ*cheV-2*, Δ*vc1384*, Δ*flgM*, and Δ*cheV-3*) were attenuated for CI (green in Fig. 2B). Of these, the Δ*flgFGHIJ* mutant produced the highest CI with a 1000-fold increased CI compared to WT, while the Δ*flgO* mutant had the lowest CI (1/1000-fold of WT). Of the other mutants, Δ*flhB*, impaired in production of the flagellar type III secretion system (fT3SS) protein FlhB, was enhanced in adaptability, and Δ*flaA*, missing the major flagellin filament protein, was attenuated. We furtherly analyzed the various phenotypes of all 26 mutants in more detail, and found that only the colonization potential in the adult mouse model differed among the mutants. All other phenotypes (e.g. motility, biofilm development, proliferation in LB/M9, H_2_O_2_ resistance, infant mice colonization) exhibited a unipolar distribution. Correlation analysis indicated that adult mice colonization was not associated with other phenotypes (Fig. S6).

**Fig. 2.**
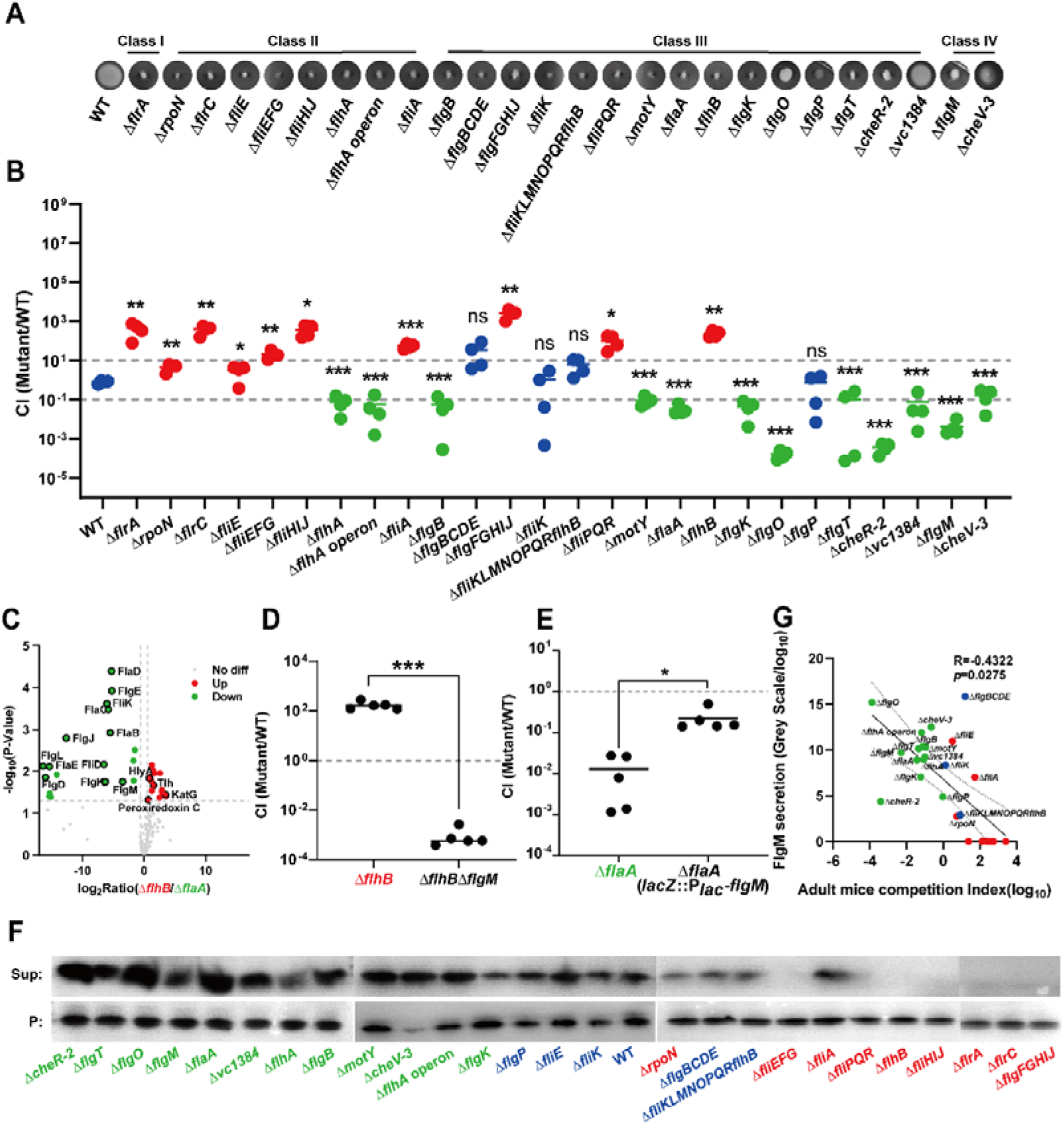
FlgM secretion regulates colonization of *V. cholerae* flagellar mutants. (**A**), Motility assay. (**B**), Competition assay in adult mice. The horizontal bar indicates the average of CI values. The CI values at 5 days post inoculation were shown. The grey dashed lines from top to bottom represent 10-fold enhanced colonization and 10-fold reduced colonization, respectively. (**C**), Volcano plot of the distribution of altered proteins. Red dots represent up-regulated expression proteins, and green dots represent down-regulated expression proteins and grey dots represent no significant expression difference (threshold > 1.5, *p* < 0.05). (**D** and **E**), Effect of FlgM on the colonization of the Δ*flhB* or Δ*flaA* mutant. The CI values at 5 days post inoculation were shown. (**F**), Western blot (WB) assay for different secretion of FlgM in the *V. cholerae* flagellar mutants. sup: supernatant, p: pellets. (**G**), Correlation analysis of FlgM secretion and CI values in adult mice model (ImageJ and Pearson method). The colors of red, green dots represent a higher, lower of colonization than WT, respectively. Significance was determined by *t*-test; *P*-values: ns, not significant, *, < 0.05, **, < 0.01, ***, < 0.001.

### FlgM secretion negatively regulating adaptability

Considering FlaA and FlhB are structural flagellar component without a regulatory function, they were confirmed through transmission electron microscopy that the 2 mutants lacked flagella (Fig. S7A), and that overexpression of the relevant genes restored their motility capability (Fig. S7B). So, we choose Δ*flhB*, Δ*flaA* and WT normalized with CFU (Fig. S7C) were to explore the underlying mechanism of these opposite colonization properties by sequenced proteins in the supernatant (Fig. 2C). These proteins from Δ*flhB* and Δ*flaA* were compared to a WT normalized control, in an attempt to seeking differentially expressed proteins. This detected 36 proteins that were differently secreted between the Δ*flhB* and Δ*flaA* mutants (Table S6, Fig. S7D), including 15 flagellar assembly proteins, 9 cellular processes related proteins, 2 secreted hemolysin proteins, 2 cell membrane associated proteins, 2 reactive oxygen species (ROS)-resistance proteins, and 6 other proteins.

Hemolysins HlyA and Tlh are secreted toxin proteins that play a role in the virulence of *V. cholerae* [6, 39]. However, their deletions in Δ*flhB* to give Δ*flhB*Δ*hlyA* and Δ*flhB*Δ*tlh*, Δ*flhB*Δ*katG* and didn’t affect their CI in the adult mice model (Fig. S8A and B). Colonization in this model can also be enhanced by ROS-resistance, for which KatG and catalase are held responsible [32]. Experiments were performed to investigate this, for which N-acetylcysteine (NAC) was employed to remove ROS from the murine intestine (Fig. S8C), but this revealed no difference in the CI of Δ*flhB* or Δ*flaA* compared to that in absence of NAC (Fig. S8D and E). As not all mutants but the majority are resistant to ROS, this did not seem to explain the enhanced colonization of particular flagellar mutants (Fig. S9).

Interestingly, the secretion of FlgM in Δ*flhB* was reduced about 9-fold compared with that of Δ*flaA* (Fig. 2C). FlgM is an anti-σ^28^ factor, and functions as a chaperone protein of σ^28^ factor FliA [40]. Our previous research had demonstrated that FlgM can be secreted by Δ*flgD* through broken flagella, which activates FliA activity, to then repress the expression of *hapR* that subsequently upregulates the expression of virulence gene *tcpA* and enhances *V. cholerae* colonization in infant mice [4]. Introduction of a double-mutation where *flgM* was knocked out in a Δ*flhB* background dramatically reversed the Δ*flhB* colonization pattern, from a CI of 176.3293 for Δ*flhB* to 0.0098 for Δ*flhB*Δ*flgM* (*P*= 0.0005) (Fig. 2D). In addition, the overexpression of FlgM in Δ*flaA* (*lacZ*::P*_lac_-flgM*) also significantly reversed the phenotype of Δ*flaA*, from CI=0.0130 for Δ*flaA* to 0.2285 for Δ*flaA* (*P*= 0.0146) (Fig. 2E). Based on these observations, we analyzed FlgM secretion in all 26 mutants and found that this negatively correlated (R^2^=-0.4322, *P=*0.0275) with their CI in adult mice (Fig. 2E and F). Together, these data suggest that changes in intracellular FlgM levels in the various flagellar mutants may contribute to their diverse colonization properties in adult mice.

### Intracellular FlgM drives hapR expression

*V. cholerae* employs cell-to-cell communication, so-called quorum sensing, of which HapR is the master regulator, which is in charge of many important regulation activities in response to different environmental signals [41, 42]. In the Δ*flhB* mutant, a high concentration of intracellular anti-σ^28^ factor FlgM is available to bind to σ^28^ factor FliA, thereby inhibiting its function; this leads to an increase in *hapR* transcription (Fig. 3A, left) [4]. We sequentially knocked out *flgM*, *fliA*, and *hapR* in a Δ*flhB* background. Whereas the double mutant Δ*flhB*Δ*flgM* had a reduced CI (0.0010) compared to Δ*flhB* (CI=210.5072, *P*=0.002), this phenotype was more than reversed in the triple mutant Δ*flhB*Δ*flgM*Δ*fliA* (CI=616.9768), whereas the quadruple mutant Δ*flhB*Δ*flgM*Δ*fliA*Δ*hapR* had a reduced CI (1.408) again (Fig. 3A, right). The order of sequential mutation was then reserved by inactivating (in that order) *flgM*, *fliA*, and *hapR* in a Δ*flaA* background, which resulted in similar switches in colonization ability (Fig. 3B). This suggests that *hapR* expression may be responsible for the different colonization abilities of these flagellar mutants in the applied adult mouse model.

**Fig. 3.**
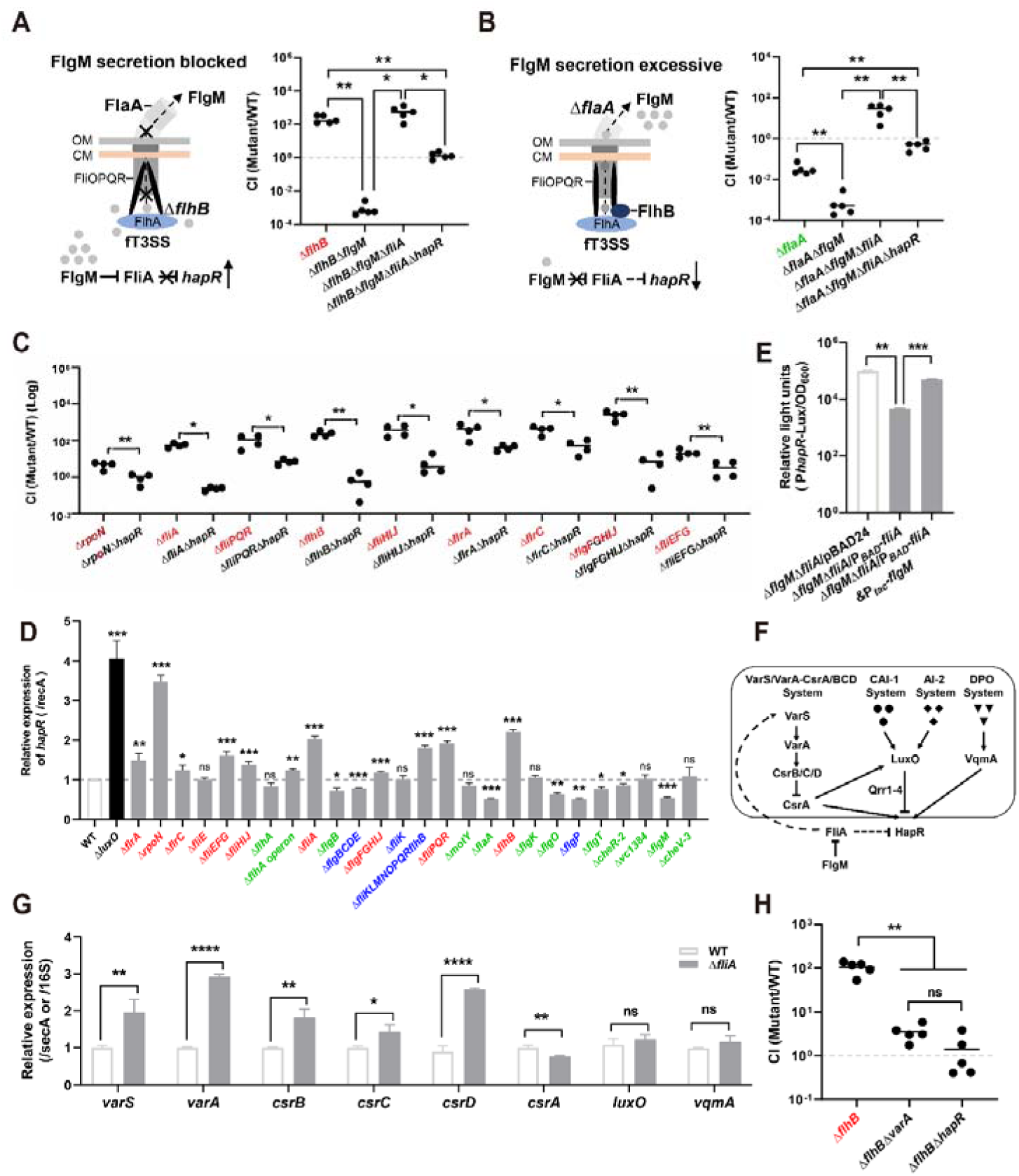
Intracellular FlgM deactivated FliA to promote *hapR* expression on transcriptionally and post-translationally. (**A** and **B**), Schematic diagram of FlgM secretion in Δ*flhB* and Δ*flaA*, and the corresponding role of FlgM-FliA-*hapR* pathway in colonization, respectively. (**C**), RT-PCR results of *hapR* expression in flagellar mutants with enhanced colonization. The *recA* was used for normalization and Δ*luxO* was used as a positive control. (**D**), *In vivo* competition experiments of different *hapR* mutants in adult mice. The CI values at 5 days post inoculation were shown. (**E**), Effect of FlgM and FliA overexpression on *hapR* expression. (**F**), Quorum sensing (QS) regulatory network of *V. cholerae*. (**G**), Gene expression in the QS regulatory pathway. Gene expression was normalized by *16S rRNA*, except for small RNAs called *csrBCD* which were normalized by *secA*. (**H**), Role of the VarS/VarA-CsrA/BCD system in adult mice colonization. The CI values at 5 days post inoculation were shown. The colors of red, green, blue dots represent a higher, lower of colonization than WT or same level, respectively. Significance was determined by *t*-test and one-way ANOVA analysis; *P*-values: *, < 0.05, **, < 0.01, ***, < 0.001, ****, < 0.0001.

This hypothesis was confirmed by determining that the transcription of *hapR* was significantly upregulated in those mutants with enhanced colonization, and significantly downregulated or unchanged in the other flagellar mutants (Fig. 3C). And In addition, the knockout of *hapR* significantly decreased the CI (Fig. 3D), suggesting that a FlgM-FliA-*hapR* pathway contributes to the colonization potential.

We found that overexpression of FliA (by means of pBAD-*fliA*) in Δ*flgM*Δ*fliA* significantly repressed *hapR* expression (*P*=0.020), while overexpression of FlgM in Δ*flgM*Δ*fliA* derepressed *hapR* transcription (*P*=0.003) (Fig. 3E). Similar results were obtained with *E. coli* DH5α (Fig. S10), suggesting that *hapR* expression may be regulated by FliA directly.

Previous studies had reported that the VarS/VarA-CsrA/BCD pathway, three autoinducer systems (CAI-1, AI-2, DPO), and VqmA can regulate *hapR* expression (Fig. 3F) [43–45]. We determined the expression of each key gene within the quorum sensing network in the Δ*fliA* mutant, which suggested that FlgM-FliA may indirectly inhibit HapR degradation through the VarS/VarA-CsrA/BCD pathway (Fig. 3G). To compare the priority of direct and indirect regulation of *hapR* by FlgM-FliA, the *varA* and *hapR* genes were individually knocked out in a Δ*flhB* background. Adult mice colonization results revealed that this drastically reduced colonization compared to Δ*flhB* (*P*=0.0026 for *varA* and *P*=0.0023 for *hapR*); no significantly difference was observed between Δ*flhB*Δ*varA* and Δ*flhB*Δ*hapR* (Fig. 3H). The double mutation of *varA* and *flhB* consistently decreased the expression level of *hapR* (Fig. S11). Taken together, it is concluded that interaction between intracellular FlgM and FliA derepresses *hapR* transcription, resulting in an enhanced colonization ability of *V. cholerae* in adult mice.

### HapR monitors methionine metabolism

We next compared precipitated protein profiles of Δ*flhB* and Δ*flaA*, using WT as a normalization control in Table S7. The cellular protein levels of flagellin proteins FlaD, methionine sulfoxide reductase MsrC, serine protease VesC, and hypothetical proteins VC1075 were upregulated (Fig. 4A). However, no significantly difference was observed among Δ*flhB* and Δ*flhB*Δ*flaD,* Δ*flhB*Δ*vesC,* Δ*flhB*Δ*vc1075* in the adult mice (Fig. S12). The MsrC was the highest enriched cellular protein in Δ*flhB* (Log_2_Ratio=3.000, *P*=0.002), relative to Δ*flaA* (Fig. 4A). Therefore, we now hypothesize that cellular methionine contributes to the different colonization patterns of the 26 flagellar mutants in adult mice.

**Fig. 4.**
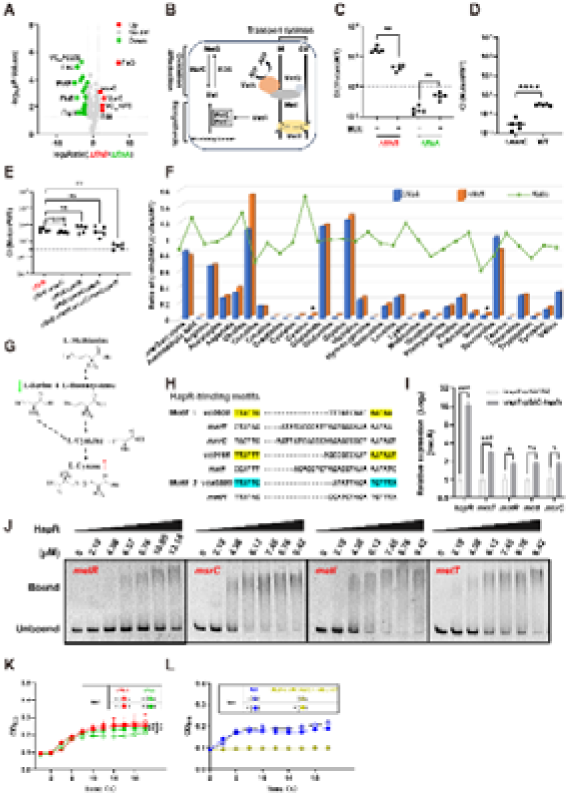
HapR regulates *V. cholerae* colonization through methionine metabolism and transportation. (**A**), Volcano map of the precipitated proteomic of Δ*flhB/*Δ*flaA*. Red dots represent up-regulated expression proteins, and green dots represent down-regulated expression proteins and grey dots represent no significant expression difference (threshold > 1.5, *p* < 0.05). (**B**), Schematic diagram of methionine (Met) metabolism. (**C**), Effect of Met addition (in drinking water) *in vivo* on the colonization of Δ*flhB* and Δ*flaA*. (**D**), Effect of *msrC* in precipitated proteomic on the colonization of Δ*flhB* mutant. (**E**), Effect of indicated mutants on the colonization of Δ*flhB,* Δ*flhB*Δ*msrC*, Δ*flhB*Δ*metR*, Δ*flhB*Δ*metR*Δ*msrC*, Δ*flhB*Δ*metR*Δ*msrC*Δ*metI*Δ*metT*. (**F**), Amino acids and its derivatives were quantified by UPHLC. The column diagram represents the contents of amino acid and its derivatives in Δ*flhB* and Δ*flaA*. The line chart represents the ratio of amino acid and its derivative content of Δ*flhB*/Δ*flaA*. (**G**), Metabolic pathway of methionine converted to other amino acids. Methionine can be converted to homocysteine by demethylation, which subsequently forms cysteine together with serine. Two molecules of cysteine synthesize cystine, while serine itself can be converted to cysteine. The metabolic pathway was annotated by Kyoto Encyclopedia of Genes and Genomes. (**H**), Promoter sequence alignment of *metR*, *msrC*, *metI*, and *metT* with conserved two HapR binding Motifs. Yellow shading represents the conserved binding sequence of Motif 1 and aqua shading represents the conserved binding sequence of Motif 2. (**I**), Effect of *hapR* overexpression on the expression of *metR*, *msrC*, *metI*, and *metT*. The bars represent the corresponding standard deviations. (**J**), EMSA for HapR binding to the promoters of *metR*, *msrC*, *metI*, and *metT*. (**K**), Growth curve of Δ*flhB* and Δ*flaA* in the MM medium with or without 0.01mM L-Met. (**L**), Growth curve of WT, Δ*flhB*Δ*metR*Δ*msrC*Δ*metI*Δ*metT* in the MM medium with or without 0.01mM L-Met. The colors of red, green, blue dots represent a higher, lower of colonization than WT or same level, respectively. The CI values at 5 days post inoculation were shown. The horizontal bars indicate the averages of CI. Significance was determined by *t*-test and one-way ANOVA methods; *P*-values: ns, not significant, *, < 0.05, **, < 0.01, ***, < 0.001.

In *V. cholerae*, cellular methionine metabolism is finely regulated via biosynthesis, transformation and transportation systems [Fig. 4B] [46]. As expected, the addition of methionine significantly reduced Δ*flhB* and it increased Δ*flaA* colonization in adult mice (Fig. 4C), whereas Δ*msrC* drastically reduced the colonization advantage (Fig. 4D, CI=0.015). However, knockout of the methionine biosynthesis genes *msrC* or/and *metR* in a Δ*flhB* background didn’t reduce colonization advantage, unless the methionine transportation genes *metI* and *metT* were also inactivated (Δ*flhB*Δ*msrC*Δ*metR*Δ*metI*Δ*metT vs.* WT compared to Δ*flhB vs.* WT, FC=6.341, *P*=0.001) (Fig. 4E). This suggests that the cellular methionine yield is precisely regulated by a range of metabolic and transportation genes in *V. cholerae*, which is not surprising, since methionine is an essential amino acid for bacterial metabolism and proliferation. Subsequently, the amino acid content in the precipitates of Δ*flaA*, Δ*flhB*, and WT were quantified. We found lower serine concentration (*P*=0.040) and higher concentrations of cystine (a disulfide of two oxidized cysteine molecules, *P*=0.018) in Δ*flhB* compared with Δ*flaA*, but no variation in methionine concentrations could be demonstrated by intracellular amino acid assays (Fig. 4F). Beyond bacterial biosynthesis and uptake, intracellular methionine can also be increased via other amino acid pathways. The KEGG pathway shows that L-cystathionine, a downstream product of serine and methionine, can be converted to cysteine, from which cystine is formed (Fig. 4G) [47], so that a higher active methionine metabolism in Δ*flhB* may relate to lower serine concentrations and higher cystine concentrations. Interestingly, we also found that *hapR* is involved in the expression of enzymes related to other amino acid metabolism by analyzing the transcriptome from our Mr.VC database [48, 49], including serine and cysteine metabolism, suggesting that *hapR* may be involved in the regulation of methionine metabolic flow.

Tsou et al. have described that HapR is a global regulator in *V. cholerae,* playing an important role in virulence regulation, biofilm development, natural transformation, and stress response [42]. By means of sequence alignments we identified sequences characteristic of the HapR-binding motif 1 in front of *metT*, *msrC*, *metI*, and a HapR-binding motif 2-like sequence in front of *metR* (Fig. 4H). Indeed, HapR directly promotes the transcription of these four genes, as demonstrated by RT-qPCR (Fig. 4I), and binding of HapR to promoter DNA was demonstrated by EMSA experiments (Fig. 4J, Fig. S13, Table S5). We observed that it rescued the growth defect of Δ*flaA* in methionine-limited medium (MM) (Fig. 4K) and adding methionine to the medium did not revive the growth of Δ*flhB*Δ*metR*Δ*msrC*Δ*metI*Δ*metT* (Fig. 4L).

### Methionine metabolism affects V. cholerae colonization

To examine the metabolic activity of Δ*flhB*, Δ*flaA* and WT in vitro, we used FITC-labeled glucose method (Fig. 5A). The metabolic activity is often labeled by marking bacterial cell wall precursors [50, 51], such as GlcNAc-1-phosphate, which is originally derived from glucose [52]. Fluorescein Isothiocyanate (FITC) is a commonly used fluorescent probe molecule that can form a covalent complex with glucose molecules. After FITC-labeled glucose is utilized, the detection of fluorescence signals can indicate bacterial metabolic activity. A transwell co-culture system with WT showed that the population of Δ*flhB* was highly metabolically active and 1.7-fold higher than WT (*P*=0.012), while Δ*flaA* was 0.6-fold lower (*P*=0.003) than WT (Fig. 5B). We also investigated bacterial metabolism in vivo by administering FITC-labeled glucose to mice that were colonized with either a Δ*flaA* and WT mixture or a Δ*flhB* and WT mixture, after which the bacterial content was isolated from their intestine and subjected to flow cytometry to sort those bacterial cells with high metabolic activity. These were enumerated on X-Gal plates, which revealed that the population of Δ*flhB* was 7-fold higher (*P*=0.017), while the population of Δ*flaA* was 0.2-fold lower compared to WT (*P*=0.001) (Fig. 5C). Thus, both in vitro co-culture and in vivo in adult mice co-colonization experiments provided strong evidence that bacterial metabolic activity contributed benefited the bacteria. In combination with the other results, this suggest that differences in methionine-related metabolic activity may contribute to the different colonization patterns of the flagellar mutants.

**Fig. 5.**
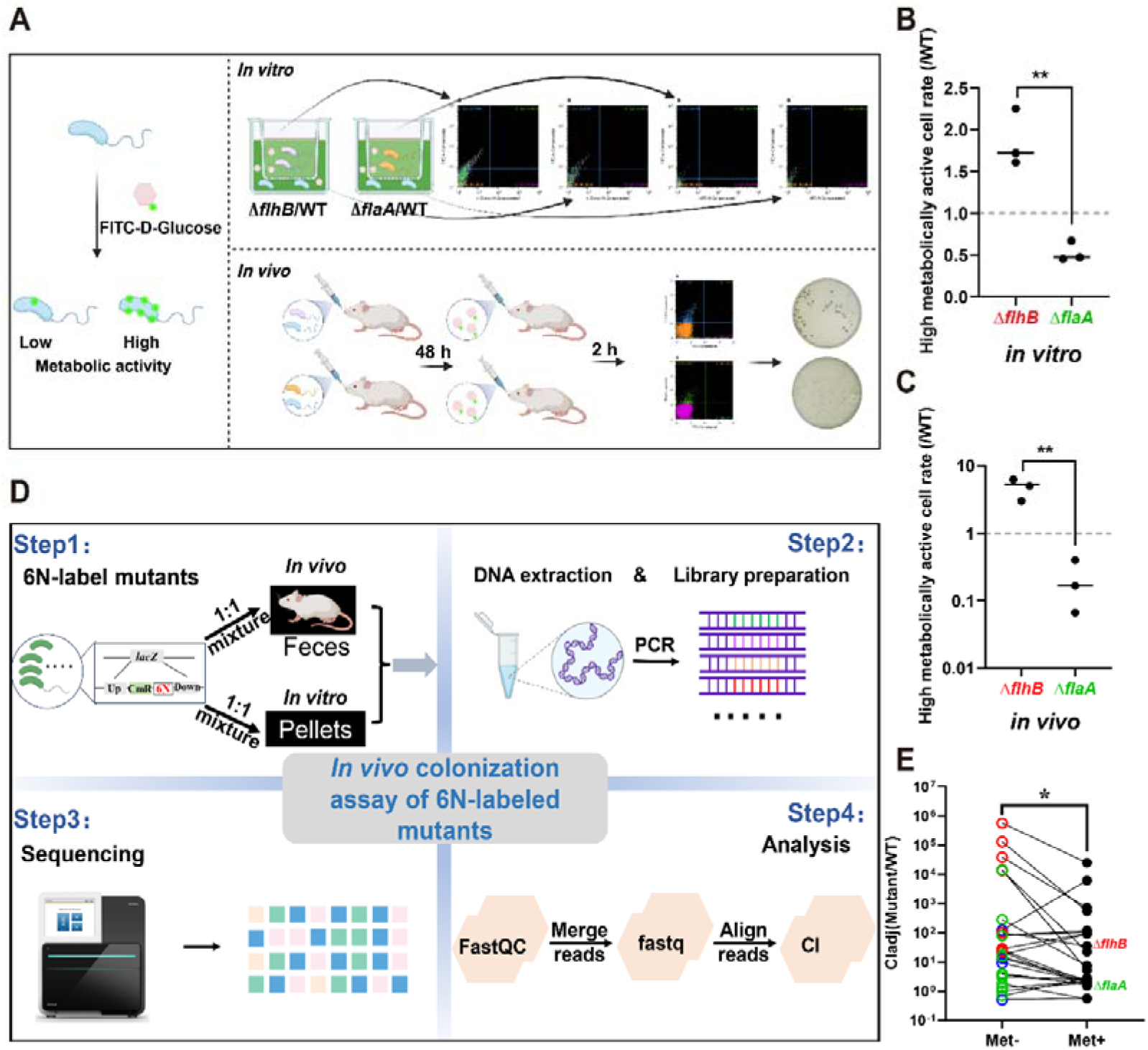
Methionine-related metabolic activity affects *V. cholerae* colonization. (**A**), The workflow for metabolic activity assay with FITC-D-Glu-labeled method in vitro and in vivo. The figure was created by BioRender. (**B**), Flow cytometry analysis of wild-type co-with Δ*flhB* and wild-type co-with Δ*flaA* FITC-D-Glu-labeled in transwell chamber at 30 min. (**C**), The bacterial metabolic activity of wild-type co-with Δ*flhB* and wild-type co-with Δ*flaA in vivo* at 48h post-inoculation. (**D**), The workflow of colonization assay of 6N-labeled mutants *in vivo*. (**E**), Colonization assay of 6N-labeled mutants *in vivo*. The raw data was analyzed by Vsearch and Python. The adjusted CIs of all flagellar mutants were analyzed by was calculated as the reads of mutant to WT colonies normalized with the input reads. The colors of red, green, blue dots represent a higher, lower of colonization than WT or same level, respectively. Significance was determined by *t*-test and one-way ANOVA methods; *P*-values: ns, not significant, *, < 0.05, **, < 0.01, ***, < 0.001.

### Ecological simulation of flagellar mutants reveals trade-off mechanisms between motility loss and host adaptability

To assess the host adaptability of the flagellar mutants, the previously characterized mutants were labeled using a novelty developed 6N-labeling method (using the primes in Table S5) and subsequently administered to adult mice via oral gavage. Colonization competition was then assessed by high-throughput sequencing (Fig. 5D, Table S8). In the absence of methionine supplementation, the competition indices (CIs) of all 26 mutants showed a scattered and diverse distribution, consistent with the head-to-head colonization outcomes presented earlier (Fig. 2B). Upon methionine addition, the distribution of CIs for all flagellar mutants became markedly more constrained, as detailed in Supplementary Dataset 6. Further pairwise analysis revealed that methionine supplementation could reverse the colonization pattern of individual flagellar mutants (Fig. 5E). Collectively, these findings demonstrate that ecological simulation of flagellar mutants reveals trade-off mechanisms between motility loss and host adaptability.

## Discussion

By employing big data analysis on *V. cholerae* life cycle samples, this study elucidates the trade-off mechanism whereby flagellar mutations result in motility loss yet enhance host adaptability through the FlgM-FliA-HapR-methionine axis, a driver of metabolic reprogramming (Fig. 6).

**Fig. 6.**
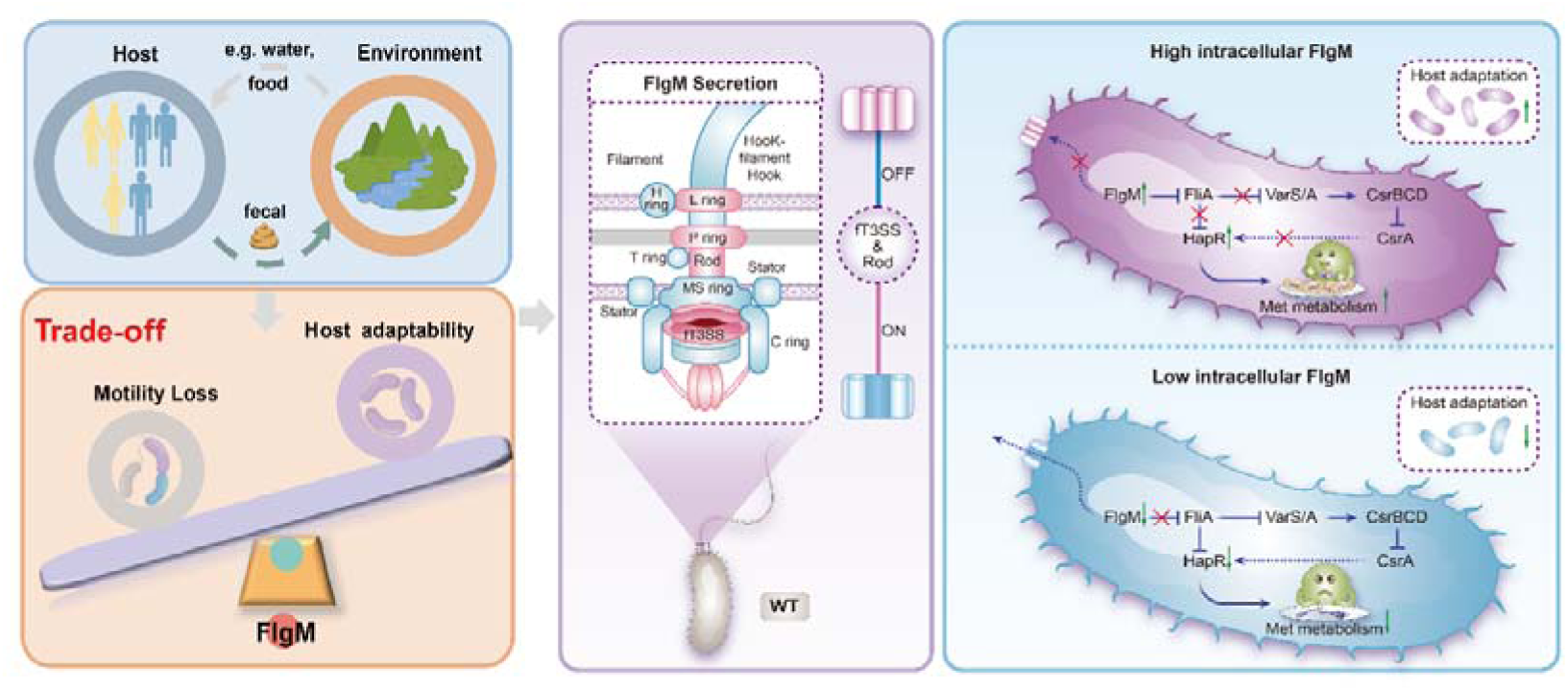
Anti-σ28 factor FlgM secretion regulates *Vibrio cholerae* adaptability in adult mice through quorum sensing and methionine metabolism. *Vibrio cholerae* predominantly cycles between aquatic reservoirs and the human host. Humans become infected by ingesting water or food contaminated with *V. cholerae*. The bacteria then colonize the small intestine, where they adapt by modulating their gene expression profile. Ultimately, they are shed back into the aquatic environment via feces, completing the transmission cycle and enabling potential new rounds of infection. During the life cycle of *Vibrio cholerae*, flagellar gene mutations are highly prevalent, clustering in genes including *flhB*, *fliA*, *fliF*, *fliD*, and *fliM*. Follow-up experimental investigations elucidated a fundamental trade-off mechanism of *Vibrio cholerae* fitness in the host gut by FlgM secretion. The level of FlgM secretion is primarily regulated by the rod complex and the flagellar type III secretion system (fT3SS). Specifically, elevated intracellular concentrations of FlgM inhibit the alternative sigma factor σ^28^ (FliA). FliA itself via the VarS/VarA-CsrA/BCD system affects the expression of quorum sensing master regulator called *hapR.* Moreover, HapR can directly bind to the promoters of genes involving in Met transportation to regulate methionine availability and metabolic activity by labeled FITC-D-Glucose to regulates *V. cholerae* colonization in the gut.

Although flagellar mutations are prevalent across the life cycle environments of *V. cholerae*, their evolutionary paths across distinct host niches remain unclear. Previous work indicates that the cyclin gene *dps*, rather than DNA repair or ROS/RNS scavenging systems, significantly increases flagellar mutation frequency in an adult mouse model [15]. In amoeba hosts, mutations in the transcriptional regulator *flrA* provide a competitive edge, while intense microbial competition in the gut may further shape *V. cholerae* evolution [16]. In addition, the *flrA* in *V. cholerae* senses intracellular levels of the second messenger cyclic diguanylate monophosphate (c-di-GMP) in response to dietary signals. Under a high c-di-GMP environment induced by casein, FlrA represses the expression of the type VI secretion system (T6SS), thereby impairing the ability of *V. cholerae* to compete with intestinal commensal bacteria via T6SS and ultimately leading to a colonization defect in casein-fed mice [53]. Furthermore, intense microbial competition in the gut may serve as a major driving force for the evolution of *V. cholerae* within the intestinal environment [54]. Integrating comparative genomics with experimental evolution assays may unravel whether these mutations represent adaptations to host niches or collateral effects of pathogen transmission dynamics.

Methionine is an essential amino acid utilized by bacteria in various physiological and metabolic processes, including polyamine synthesis, DNA synthesis, redox balance, and DNA and histone methylation [55]. Also, it is important to note that methionine plays an essential role in the immune system through its metabolites. For example, in response to antigens, T cells activate the methionine cycle to support proliferation and differentiation in cancer, indicating the importance of the methionine cycle to T cell immunity [56]. Methionine supplementation or restriction can intervene in the natural antioxidant capacity of an organism by leading to the production of endogenous enzymes that reduce oxidative stress and DNA damage [57]. These data imply that *V. cholerae* may employ the methionine pathway as a means of immune evasion. To successfully propagate and cause disease, *V. cholerae* also evades the immune system by repressing the mannose-sensitive hemagglutinin pilus via the transcription factor ToxT, preventing secretory IgA binding and enabling intestinal colonization to bypass host defenses [58].

Methionine metabolism is a central node within the interconnected amino acid network of *V. cholerae* [59]. Beyond its metabolic functions, methionine serves as a precursor for S-adenosylmethionine (SAM), a cofactor involved in critical processes including the synthesis of the quorum-sensing autoinducer CAI-1 [60,61]. Similarly, threonine, another methionine-derived metabolite, may contribute to autoinducer DPO synthesis, and in *Pseudomonas aeruginosa*, SAM participates in producing AI-1, AI-2, and AHLs [62]. Analyzing methionine flux could therefore help identify key metabolites that enhance colonization in flagellar mutants, and future work should examine whether intermediate methionine metabolites or other amino acids influence *V. cholerae* colonization through bacterial or host metabolic remodeling.

This study found that HapR binds to promoters of methionine metabolism-related genes (*metR*, *msrC*, *metI*, *metT*), a result not reported in prior systematic screens of HapR targets. The discrepancy may stem from limitations in bioinformatic prediction accuracy and variability in HapR-binding sequences [42]. Beyond methionine regulation, HapR suppresses toxin-coregulated pilus (TCP) and cholera toxin (CTX) genes by repressing *aphA*, while activating extracellular proteases and modulating T6SS genes [63–65]. However, in adult and infant mouse models, this virulence factors do not substantially affect colonization [39,66]. HapR also inhibits biofilm formation via cyclic di-GMP modulation and *vpsT* repression [67–69], although flagellar mutants show enhanced biofilm capacity, they still exhibit colonization defects, indicating biofilm is not a major driver of colonization in these strains. Additionally, HapR and RpoS jointly facilitate stress adaptation, luminal dispersal, and host exit. Although the flagellar mutants of *V. cholerae* exhibit enhanced biofilm formation capacity, they still display a colonization defect (Fig. S3, Fig. 2B), suggesting that biofilm formation is also not a major factor contributing to enhanced colonization in these mutants. In addition, HapR and RpoS collectively facilitates stress adaptation to host factors, dispersal within the intestinal lumen, and eventual exit of *V. cholerae* from the host [70]. In the precipitated proteome, we observed that MsrC was the most highly enriched cellular protein in Δ*flhB* (Log2Ratio = 3.000, P = 0.002) relative to Δ*flaA* (Fig. 4A). Through experimental validation, we concluded that methionine contributes to the differential colonization patterns among the 26 flagellar mutants in adult mice.

The flagellum renders bacteria motility, but it is also a major antigen recognized by the host [71, 72]. Extensive studies have shown that flagellar mutants of *V. cholerae* are frequently present in clinical and environmental samples, and specifically the *V. cholerae* Haiti strains contain multiple mutational hotspots in flagellum gene regions according to SNP analysis [11]. Besides mutations, *V. cholerae* also actively loses flagellar filaments during environmental changes such as mucus penetration [4], biofilm development [10], and host Lypd8 promotes the segregation antibody capture [72]. This is also demonstrated for other pathogenic bacteria. The freshwater bacterium *Caulobacter crescentus* loses its flagella through ejection trigged by ClpA [73]. The loss of the polar flagellum in *V. fischeri* and in plant-associated *Methylobacteria* has also been reported [74, 75]. Bacterial flagella also detach during starvation [76].

We demonstrated that the trade-off mechanism between motility loss and host adaptability operates via the sophisticated FlgM-FliA-HapR-methionine metabolic reprogramming axis. However, this study has the following limitations: (1) The mechanistic focus on the FlgM-FliA-HapR-methionine axis, though insightful, may not capture the full complexity of the flagellar regulatory network, and deeper metabolic validation is needed; (2) It does not directly simulate or verify how this trade-off process dynamically occurs and becomes fixed within a population under environmental pressure, lacking direct experimental evolution evidence in human; (3) The adaptive efficacy and stability of the revealed mechanism in such highly complex real-world environments require further evaluation; (4) Time-course experiments were not provided to track the changes in FlgM secretion, HapR expression, and methionine metabolism over the course of infection.

## 5. Conclusion

Based on extensive sample analysis, this study reveals that flagellar mutations in *V. cholerae* cluster in key structural and regulatory genes, underscoring their central role in host adaptation. Specifically, the secretion level of the flagellar regulator FlgM drives a phenotypic shift from motility to host adaptability. This transition is mediated through the FlgM-FliA-HapR-methionine axis, which balances the loss of motility with enhanced host adaptability. Ultimately, impaired motility promotes environmental fitness by activating quorum sensing and reprogramming methionine metabolism, elucidating a novel evolutionary trade-off strategy between movement and adaptability at the molecular level.

## Supporting information

The supplementary tables will be used for the link to the file on the preprint site.

The supplementary material will be used for the link to the file on the preprint site.

Ethical Statement

Permit Number

## Credit Authors Statement

**Guozhong Chen:** Conceptualization, Methodology, Investigation, Project administration, Funding acquisition, Visualization, Writing-original draft, Writing-review & editing. **Zixin Qin:** Methodology, Investigation, Visualization. **Fenxia Fan:** Investigation, Writing-review & editing, Funding acquisition. **Mei Luo:** Writing-review & editing. **Hongou Wang:** Methodology. **Baoshuai Xue:** Methodology. **Shucheng Li:** Methodology. **Shiyong Chen:** Supervision. **Xiaoman Yang:** Investigation. **Xiang Mao:** Investigation. **Liwen Yi:** Investigation. **Chunrong Yi:** Investigation. **Wei Li:** Investigation, Supervision. **Xiaoyun Liu:** Project administration, Funding acquisition. **Biao Kan:** Supervision, Project administration. **Zhi Liu:** Conceptualization, Methodology, Visualization, Project administration, Funding acquisition, Supervision, Writing-original draft, Writing-review & editing.

## Acknowledgments

We are grateful to all the members of our team. We thank Core Facilities for Life Science, Huazhong University of Science and Technology for the technical support.

## Ethics statement

Only publicly available cholera data from NCBI database were used in this study. Informed consent was obtained from all subjects in the original studies. For those datasets, ethical review and approval can be accessed in the original studies.

All animal experiments were carried out in strict accordance with the animal protocols that were approved by the Ethical Committee of Huazhong University of Science and Technology (Approval no. 2017-S560).

## Supplementary Material

Supplementary material is available online.

## Data Availability

The proteomic data that support the findings of this study are openly available in the iProX database at https://www.iprox.cn//page/SCV017.html?query=IPX0005998000, accession number IPX0005998000. The raw data of 6N-Label is openly available in the NCBI database at https://www.ncbi.nlm.nih.gov/sra/?term=PRJNA1099378, reference number PRJNA1099378.

## Table legends

**Table S1. Sample collection.** Samples were collected from 19 countries. Based on their sources, the cholera-related metagenomic sequencing samples obtained from the SRA database were classified into five categories: human-derived, clinical, aquatic (including water and sewage), fish-derived, and amoeba-derived.

**Table S2. Structure and function of *Vibrio cholerae* flagella.** The flagella of *V. cholerae* are divided into many flagellar components which play a different role in flagellar biosynthesis. Their functions were summarized by literature search, and literature sources were added.

**Table S3. In silico analysis of flagellar gene mutations in *V. cholerae***. Mutations in flagellar genes were analyzed using the Diamond software.

**Table S4. Strains and public Plasmids used in this study.** The strains or public plasmids were descripted. The public plasmids were obtained from anywhere.

**Table S5. The primers used in this study.** These primers involve plasmid construction, Reverse Transcription Polymerase Chain Reaction, Electrophoretic Mobility Shift Assay, *lacZ* sequencing.

**Table S6. Altered proteins in secretory proteomics data.** The up-regulated and down-regulated expression proteins of Δ*flhB*/Δ*flaA* were list (Fold change > 1.5, *p* < 0.05, *t-test*). The involved functions of these altered proteins were explained.

**Table S7. Differentially expressed proteins in precipitated proteomics data.** Bacterial proteome samples of Δ*flhB* or Δ*flaA* were analyzed in triplicates. Proteins with Fold changes >1.5 and *p*< 0.05 (*t-test*) were considered as significant differences.

**Table S8. Relative abundance analysis of 6N-labelled LacZ sequence of *V. cholerae* wild-type and flagellar regulon mutants’ libraries.** The sequenced raw data was analyzed by Vsearch for merging paired-end reads and Python to calculate the proportion of reads from the 6N-labeled mutants among the total reads.

