## Supplementary material for "Anti-σ^28^ factor FlgM secretion regulates *Vibrio cholerae* adaptability in adult mice through quorum sensing and methionine metabolism": Ethical Statement

地址：武汉市航空路 13 号　　邮编：430030　　电话和传真：027-83692751

### 华中科技大学实验动物伦理委员会动物伦理审批报告

[2017]伦审字（S560）号

| | | | |
|---|---|---|---|
| 项目名称 | 霍乱弧菌和肠道微生物信号交流促进其抗逆和定殖的的分子机理 | | |
| 申请单位 | 华中科技大学生命科学与技术学院 | | |
| 项目负责人 | 刘智 | 职称 | 教授 |
| 报送资料 | 课题研究方案 | 有☑ | 无☐ |
| | 观察记录表 | 有☑ | 无☐ |
| | 研究人员名单 | 有☑ | 无☐ |
| 审查 | 研究者资格 | 符合要求☐ | 不符合要求☐ |
| | 课题研究方案 | 适当☑ | 不适当☐ |
| 有效期 | 年____月____日至______年____月____日 | | |
| 华中科技大学实验动物中心动物实验审查者意见（签名） | 报批资料审阅，符合动物伦理要求，建议按[illegible]方案！实验结束后应核查执行情况！ [illegible] 22/2-2017 | | |
| 动物实验申请者承诺在实验期间按照伦理要求进行动物实验（签名） | [illegible]　余泽年 | | |
| 动物实验申请者导师意见（签名） | 同意。　刘智 | | |

审评意见：

本伦理委员会审阅并讨论了上述相关资料，该课题研究符合《湖北省实验动物管理条例》和《华中科技大学同济医学院实验动物伦理委员会章程》，经伦理委员会审核，同意该课题实施。

华中科技大学同济医学院实验动物伦理委员会

主任签名：[illegible]

批准日期：　　年　　月　　日
